# Efficient gene knockout and genetic interactions: the IN4MER CRISPR/Cas12a multiplex knockout platform

**DOI:** 10.1101/2023.01.03.522655

**Authors:** Nazanin Esmaeili Anvar, Chenchu Lin, Xingdi Ma, Lori L. Wilson, Ryan Steger, Annabel K. Sangree, Medina Colic, Sidney H. Wang, John G. Doench, Traver Hart

**Affiliations:** Department of Bioinformatics and Computational Biology, The University of Texas MD Anderson Cancer Center, Houston, TX, USA; Graduate School of Biomedical Sciences, The University of Texas MD Anderson Cancer Center UTHealth, Houston, TX, USA; Genetic Perturbation Platform, Broad Institute of MIT and Harvard, Cambridge, MA, USA; Center for Human Genetics, The Brown foundation Institute of Molecular Medicine, The University of Texas Health Science Center at Houston, Houston, TX, USA; Department of Cancer Biology, The University of Texas MD Anderson Cancer Center, Houston, TX, USA

## Abstract

Genetic interactions mediate the emergence of phenotype from genotype, but initial technologies for combinatorial genetic perturbation in mammalian cells suffer from inefficiency and are challenging to scale. Recent focus on paralog synthetic lethality in cancer cells offers an opportunity to evaluate different approaches and improve on the state of the art. Here we report a meta-analysis of CRISPR genetic interactions screens, identifying a candidate set of background-independent paralog synthetic lethals, and find that the Cas12a platform provides superior sensitivity and assay replicability. We demonstrate that Cas12a can independently target up to four genes from a single guide array, and we build on this knowledge by constructing a genome-scale library that expresses arrays of four guides per clone, a platform we call ‘in4mer’. Our genome-scale human library, with only 49k clones, is substantially smaller than a typical CRISPR/Cas9 monogenic library while also targeting more than four thousand paralog pairs, triples, and quads. Proof of concept screens in four cell lines demonstrate discrimination of core and context-dependent essential genes similar to that of state-of-the-art CRISPR/Cas9 libraries, as well as detection of synthetic lethal and masking/buffering genetic interactions between paralogs of various family sizes, a capability not offered by any extant library. Importantly, the in4mer platform offers a fivefold reduction in the number of clones required to assay genetic interactions, dramatically improving the cost and effort required for these studies.

## Introduction

Pooled library CRISPR screens have revolutionized mammalian functional genomics. DepMap teams have screened over a thousand cancer cell lines with CRISPR knockout libraries to identify background-specific genetic vulnerabilities^1,2^, while dozens of genetic modifier screens with small molecules have explored biomarkers and mechanisms of drug sensitivity and resistance^3–9^. However, initial efforts to assay genetic interactions (GIs) – that is, the manipulation of multiple genes in the same cell to identify nonlinear combinatorial phenotypes – have proven complex and costly^10–14^. One class of GIs that has received special attention is the synthetic lethal relationships between paralogs, gene pairs or families that arise through duplication of a single ancestral gene. Functional buffering by paralogs, resulting in phenotypic masking in single gene perturbation experiments, has been shown extensively in model organisms^15,16^. Paralogs are therefore attractive targets for genetic interaction studies in human cells, and they are more easily nominated by computational analyses^17,18^ compared to genes that work in parallel pathways, such as *BRCA1* and *PARP1*^19^. Further, because the mechanism of action of drugs often relies on inhibition of paralog gene products to mediate cell toxicity, monogenic knockout in CRISPR screens have resulted in false negatives, such as the failure to identify MEK and ERK proteins as critical for cancer cell growth.

Recently, paralog synthetic lethals have been assessed with multiple CRISPR-based methods, some relying on a single Cas protein with multiple guides with others using orthogonal Cas proteins in the same cell^18,20–23^. However, with the application of different experimental and informatic pipelines, comparison across studies has been difficult. Notably, no widely accepted gold standard set of paralog synthetic lethals exists against which researchers can calculate traditional metrics of accuracy such as sensitivity and specificity. Here we describe a meta-analysis of paralog genetic interaction screens in human cells, identifying a set of background-independent paralog synthetic lethals, and demonstrate that the enhanced version of Cas12a^24^ from *Acidaminococcus sp*. (enAsCas12a^25^, hereafter referred to as Cas12a) provides the best combination of sensitivity and simplicity for genetic interaction studies.

Building on our prior work^18,26,27^, we further extend the capabilities of the Cas12a system. We show that it can reliably utilize arrays encoding four optimized gRNAs when delivered by lentivirus in a pooled screening format. We describe the in4mer platform that uses oligonucleotide synthesis to construct libraries of four-guide arrays that target specified sets of one to four genes independently.

From the in4mer platform, we develop a genome-scale human library named Inzolia which, with ∼49,000 clones, is approximately 30% smaller than standard whole-genome libraries and has the unique added capability of targeting more than 4,000 paralog families of size two, three, and four. This combination of features is not available with any other CRISPR perturbation platform.

## Results & Discussion

With the discovery that paralogs are both systematically underrepresented as hits in pooled library screens and likely offer the highest density of genetic interactions, several independent studies have each targeted hundreds of paralog pairs in multiple cell lines^18,20–23^. However, evaluating the quality and consistency of these studies has proven difficult, since each uses a different technology and custom analytics pipeline for hit calling, and overlap between the targeted paralog pairs in each study is surprisingly slim (Figure 1A,B).

**Figure 1.**
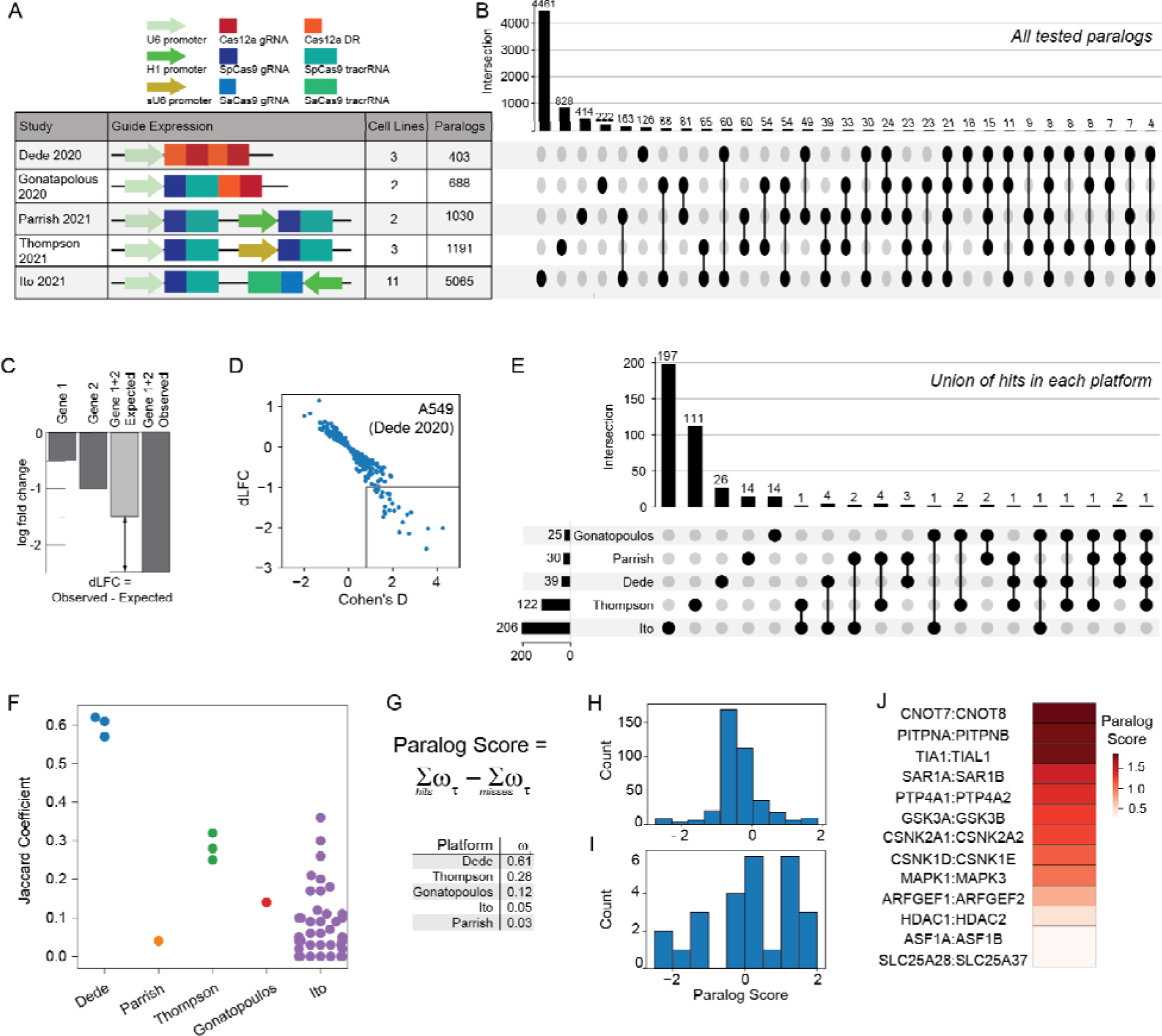
Comparative analysis of synthetic lethality screens. **A)** Different multiplex CRISPR perturbation methods applied to assay paralog synthetic lethality. B) Tested paralog pairs in each study. Upset plot shows the intersection of pairs across different studies. C) Quantifying synthetic lethality between paralog pairs. Single mutant fitness (SMF) is the mean log fold change of gRNAs that target an individual gene. Expected double mutant fitness (DMF) is calculated as the sum of SMF of gene 1 and gene 2. Delta Log Fold Change (dLFC) is the difference between observed and expected fold change and is used as a measure of genetic interaction. D) dLFC vs. Cohen’s D in one data se, A549 screen in Dede ^9^ . E) Comparison of union of hits across all cell lines in each study. F) Jaccard coefficient comparing hits across all pairs of cell lines within each study. G) The “paralog score” is t e weighted sum of hits minus the weighted sum of misses; i.e. where the gene pair is assayed but not a hit. Weights are the median of the platform-level Jaccard coefficients from (F). H). Histogram of paralog scores of 388 hits across all 5 studies. I) Histogram of paralog scores across 26 hits in >1 study. J) Thirteen candidate “paralog gold standards” with paralog score > 0.25 and hit in more than one study.

We developed a unified genetic interaction calling pipeline, based on measuring a pairwise gene knockout’s deviation from expected phenotype (delta log fold change, dLFC) and the standardized effect size of this deviation (Cohen’s d) (Figure 1C,D; Supplementary Figure 1). After performing background-specific normalization (see Methods), we classified paralogs as synthetic lethal if they met both dLFC and Cohen’s d thresholds (Methods). A total of 388 gene pairs were scored as hits across all five multiplex perturbation platforms (Figure 1E).

Using this pipeline, we found the large majority of paralog synthetic lethals to be platform-specific. To aid in comparing hits within and across experiments, we developed a platform quality score that broadly measures the replicability of these synthetic lethal screening technologies across different cell lines. We reasoned that, like individual essential genes, a large fraction of paralogs should show consistent synthetic lethality across most or all cell lines, which should be reflected in similarity of synthetic lethality profiles across cell lines. We therefore calculated the Jaccard coefficient of each pair of cell lines screened by a particular platform, then took the median of each platform’s Jaccard coefficients as the platform quality score (Figure 1F).

We then calculated a paralog confidence score for each gene pair by taking the sum of each hit, weighted by the platform quality score, and subtracting the sum of each experiment in which the pair was assayed but not deemed a hit (a “miss”), also weighted by quality score (Figure 1G). Using this approach, paralog pairs that are hits in multiple high-quality screens outweigh pairs that are hits in screens with lower replicability or pairs that are background-specific hits in high-scoring screens. We further filtered for hits that are detected by more than one platform, minimizing the bias toward paralog pairs that are only assayed in one set of screens or with one perturbation technology. We identified a total of 26 gene pairs that meet these criteria, and we classified the top 13 hits (with paralog score > 0.25) as candidate paralog synthetic lethal gold standards (Figure 1H-J). Measuring the recall of each of the 21 cell line screens against these gold standards confirmed that the Cas12a platform used in Dede et al^18^, with two Cas12a guide RNAs expressed from the same promoter, yielded the highest within-platform replicability (Supplementary Figure 2). Other platforms often showed high sensitivity in one screen, but highly variable sensitivity across multiple screens (Supplementary Figure 2).

### Optimizing the Cas12a system for multiplex perturbations

Based on the consistency of the Cas12a results in the paralog screens and its potential applications to higher-order multiplexing, we explored whether crRNA arrays longer than two guides could be utilized at scale. The Cas12a system has previously been shown to mediate multiplexing beyond two targets^28,29,22^ with varying levels of efficiency. Guide RNA design is a critical factor in all CRISPR applications^30^, and empirical data on Cas12a guide efficacy is relatively sparse compared to >1,000 whole-genome screens in cancer cell lines performed with Cas9 libraries. We tested more than 1,000 crRNA from the CRISPick^26,31^ design tool in a pooled library targeting coding exons of known essential genes and found very strong concordance between the CRISPick on-target score and the fold change induced by the gRNA (Supplementary Figure 3). We therefore considered CRISPick designs for all subsequent work.

We have previously shown that arrays encoding two crRNAs show minimal position effects^18,26,27^, but information about longer arrays is sparse^22,28,29^. We constructed arrays of up to 7 gRNAs to evaluate the maximum length that would yield gene knockout efficiency sufficient for pooled library negative selection screening. A set of seven essential and nonessential genes were selected and each assigned to one position (1-7) on the array. A single guide RNA was selected for each gene, and arrays were constructed such that all combinations of essential and nonessential gRNA were represented, for a total library diversity of 128 array sequences (Figure 2A). The process was repeated two more times, using different gRNA targeting the same genes, creating three pools each of 128 unique sequences, each targeting the same seven essential and nonessential genes in all combinations (Figure 2B).

**Figure 2.**
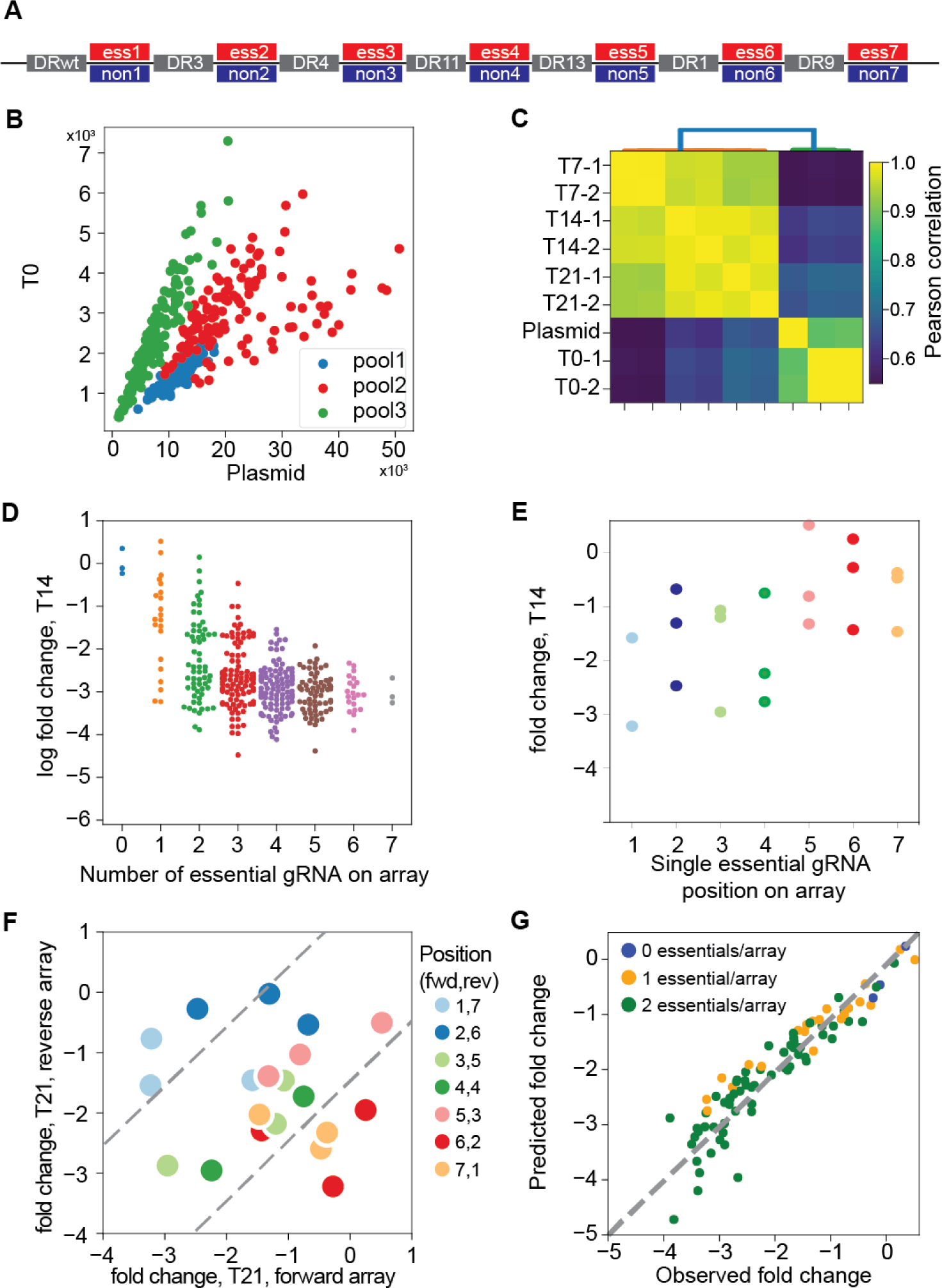
Multiplexing beyond 2 guides with Cas12a. (A) 7mer arrays were constructed with all combinations of either an essential or nonessential guide at each position (2^7=128 species), using the same DR sequences at each position, in three independent sets with unique gRNA sequences targeting the same genes at each position (n=384 total). (B) Guide sets were evenly represented in the combined pool before and after packaging and transduction (C) 7mer guide array representation is consistent across replicates and variation is consistent with high quality screens. (D) Fold change of guide arrays vs. number of essential guides on the array (n=384 arrays). (E) Fold change vs. position of essential guide on array, for all arrays encoding one essential guide (six nonessentials). (F) Fold change of guide arrays encoding one essential per array, forward vs. reverse orientation. Essential guides expressed at positions 6 and 7 deviate from the diagonal, indicating position-specific loss of editing. (G) A linear regression model that learns single gene knockout effects can be used to predict combinatorial target phenotype, an accurate null model for genetic interactions.

We cloned the 7mer pools into the pRDA_550 vector, a one-component lentiviral vector expressing the Cas12a CRISPR endonuclease and the *pac* puromycin resistance gene from an EF-1α promoter and the array of Cas12a guides from a human U6 promoter (see Methods). We used the library to screen K562 cells, a *BCR*-*ABL* chronic myeloid leukemia cell line commonly used for functional genomics, and collecting samples at 7, 14, and 21 days (Figure 2C). After normalization (see Methods), arrays with no essential gRNA showed no sign of negative selection compared to arrays with any number of essential gRNA. Arrays with multiple essential guides showed increasing loss of fitness, reaching maximal phenotype at 4 to 5 essential guides per array (Figure 2D; Supplementary Figure 4).

To evaluate position-level effects, we considered arrays encoding a single essential gRNA at any of the seven positions. Across the three replicates, we consistently observed greater fold change at the first four positions compared to the last three positions on the array (Figure 2E). We further tested whether this efficiency drop at the end of the array was a position-dependent effect or the result of unfortunate guide or gene selection. We constructed a second array with the same gRNA targeting the same genes in reverse order (one essential gRNA per array) and re-screened the same cells. When comparing the fold change of the forward array with the reverse array, observed fold changes on the diagonal indicate gene- and guide-level effects independent of position, while deviations from the diagonal indicate position-specific effects. Our data confirm that the first four to five gRNAs show no position-specific effects, but positions six and seven show marked deviation from the diagonal (Figure 2F). Based on these observations, we conservatively conclude that the Cas12a system using the pRDA_550 vector can effectively express and utilize arrays of four gRNAs.

We also evaluated whether the 7mer array could be used to identify combinatorial phenotypes.

We trained a linear regression model using a binary encoding of guide arrays as a predictor (where nonessential = 0 and essential = 1) and log fold change as a response variable (see Methods). The regression model provides excellent prediction of fold change for arrays encoding two essentials (R^2^ = 0.78-0.91 for the three pools) from the sum of calculated single-guide position-level regression coefficients (Figure 2G). These observations are consistent with the multiplicative model of genetic interactions, which predicts that the result of independent loss of fitness perturbations is the sum (in log space) of the fold changes of the individual fitness perturbations. It further supports the utility of the Cas12a platform for multiplex perturbation and detection of genetic interactions, which are simply deviations from the expected phenotype according to this model, because the null model accurately fits the data for independent combinatorial perturbations.

### The in4mer platform for single and combinatorial perturbation

With confidence that the Cas12a platform supports independent utilization of four guides expressed from a single array, we designed a prototype genome-scale library that targets both protein-coding genes and paralog families in the same pool. Each array encodes four distinct gRNA, each with a different DR sequence selected from the top performers in DeWeirdt *et al*^26^. (Figure 3A). The library targets each of 19,687 protein coding genes with one four-guide array encoding 20mer crRNA sequences from the top four guides nominated by the CRISPick algorithm, and a second four-guide array using the same guides in a different order. The library also targets 2,082 paralog pairs with a single array encoding two gRNA per gene and a second array encoding the same gRNA in a different order (see Methods and Supplementary Figure 5 for paralog selection strategy). Additionally, 167 paralog triples and 48 paralog quads are targeted by two arrays, with each array encoding a single guide targeting each gene. Arrays targeting triples are padded with a fourth guide targeting a randomly selected nonessential gene. For triples and quads, the two arrays per set encode different gRNA sequences (Figure 3A). Total library size is 43,972 4mer CRISPR arrays, including 4 arrays with 4 guides each targeting EGFP. Since the leading direct repeat sequence is already on the pRDA_550 backbone, the library can be synthesized as a 208mer oligo pool, including 5’ and 3’ amplification and cloning sequences (see Methods).

**Figure 3.**
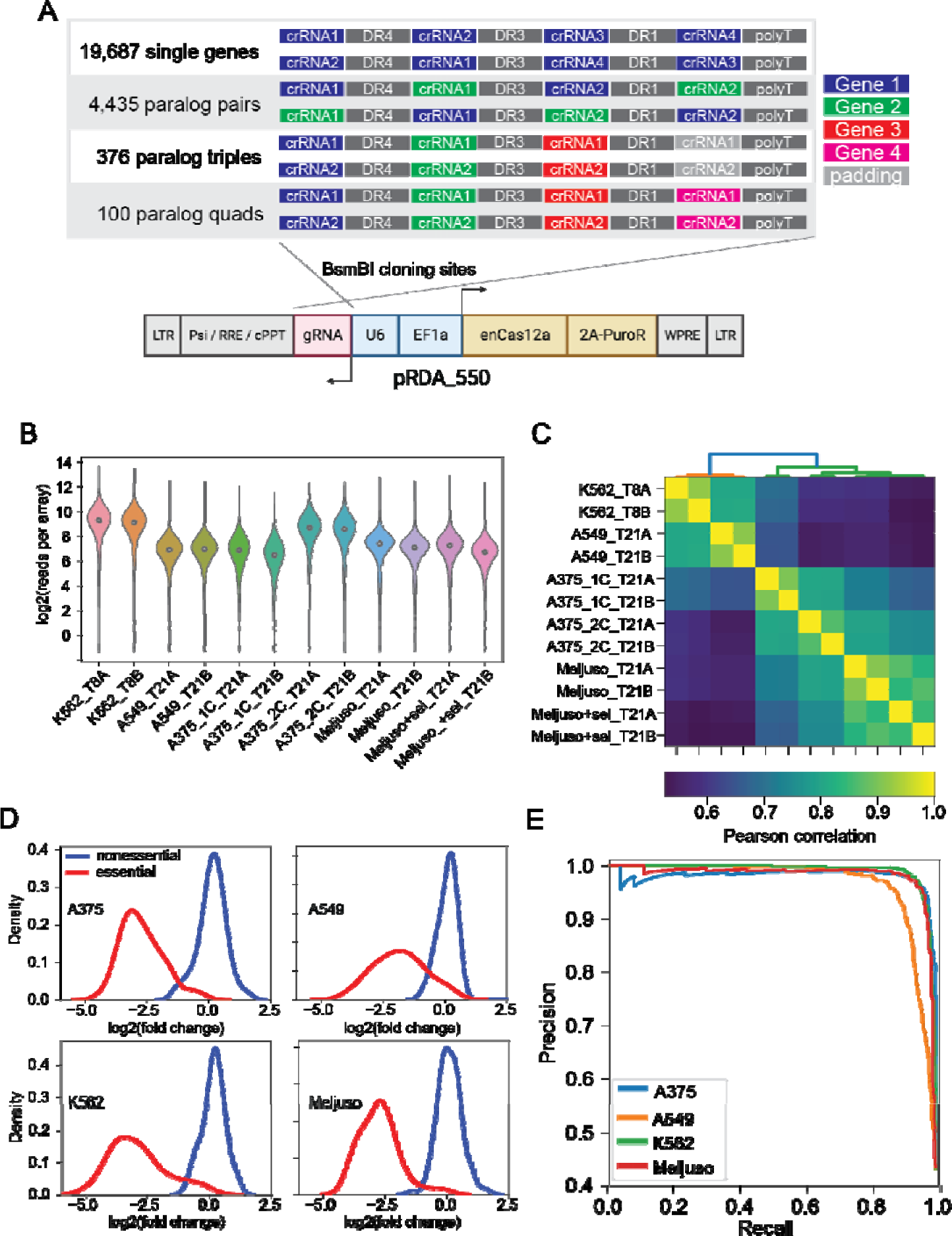
In4mer platform for whole-genome screening. (A) Inzolia human whole-genome library targets single genes and paralog pairs, triples, and quads with arrays of 4 Cas12a gRNA. Each gene or gene family is targeted by two arrays encoding the same gRNA in different order. Commercially synthesized oligo pools are cloned into the one component pRDA_550 lentiviral vector; schematic created in Biorender. (B-F) Screening in K562 CML cells and A459 lung cancer cells. (B) Read cou ts from the plasmid and experimental timepoints after lentiviral transduction. (C) Correlation of sample read counts. Endpoint replicates are highly correlated. (D) Fold change distributions of arrays targeting reference essential (red) and nonessential genes (blue) in four cell lines. (E) Precision/recall analysis from ranked mean fold change of arrays targeting each gene, calculated against reference essential and nonessential genes.

We conducted screens in K562, and in A549, a *KRAS* lung cancer cell line with wildtype *TP53*, using standard CRISPR screening protocols (500x library coverage, 8-10 doublings). Array amplicons were sequenced using single-end 150-base Illumina sequencing. Quality control metrics met expectations (Figure 3B), and the abundance distributions of endpoint replicates were highly correlated (Figure 3C). Fold changes showed increasing correlation when comparing clones targeting the same individual gene or paralog family within one replicate (n=22k targets, r=0.78); all clones between two technical replicates derived from the same transduction (n=44k arrays, r=0.86); and the mean of clones targeting the same gene across technical replicates (n=22k targets, r=0.92) (Supplementary Figure 6). In4mer guide arrays effectively discriminated reference essential genes from nonessentials (Figure 3D), yielding precision-recall curves comparable to other CRISPR/Cas9 and Cas12a whole genome screens (Supplementary Figure 7).

Building on the success of the prototype library, we designed a second human genome library with several modifications. The new library targets roughly twice as many paralogs (4,435 pairs, 376 triples, and 100 quads; Figure 3A and Supplementary Figure 5), includes nontargeting arrays to facilitate the estimation of fitness effects arising from multiplex locus-nonspecific DNA double strand breaks, and other minor technical changes such as using nontargeting sequences to extend 3mer paralog constructs into 4mer guide arrays, instead of random selection of nonessential guides as used in the prototype. This library, which we call Inzolia, contains ∼49k unique arrays (Figure 3A), and was cloned into both the pRDA_550 one-component vector and pRDA_052 guide-only expression vector for two-component CRISPR/Cas12a systems.

We screened the Inzolia library in Meljuso melanoma cells with the two-component (split-vector) library and A375 melanoma cells with both the one- and two-component systems. Both cell lines effectively identified essential and nonessential genes (Figure 3D,E) and the screens showed results consistent with previous Cas12a screens using the Humagne library (Supplementary Figure 7). Further, the one-component (pRDA_550) and two-component (pRDA_174 + pRDA_052) libraries yielded equivalent results (Supplementary Figure 7).

The whole-genome libraries target small paralog families as well as single genes. To evaluate paralog genetic interactions, we used the multiplicative model to calculate the expected fitness of pairwise knockouts by summing the log fold change of the single gene knockouts (Figure 1C). We then compared the observed mean fold change of guide arrays targeting gene pairs with the expected fold change under the multiplicative null model to calculate a dLFC that represents the magnitude of the genetic interaction (Figure 4A). Gene pairs with strongly negative dLFC are highly concordant with the gold standard paralog synthetic lethals described above. The Inzolia library targets 24 of the 26 gene pairs that are hits in >1 of the previously published screens, and 12 of the 13 candidate gold standard paralog synthetic lethals with paralog scores >= 0.25. Of those 12, 9 have dLFC < -1 in Meljuso cells, for an estimated sensitivity of 75% (Figure 4B). Moreover, all 12 pairs (100%) with high paralog score are essential, regardless of synthetic lethality, as are 10 of 12 pairs (83%) with lower paralog scores (Figure 4C), consistent with either synthetic lethality or one paralog being essential. Many other paralogs show genetic interactions as strong as these positive controls (see Figure 4D for selected examples), with sequence similarity between paralogs being a strong predictor of GI (Figure 4E), in keeping with prior observations by De Kegel & Ryan^17^.

**Figure 4.**
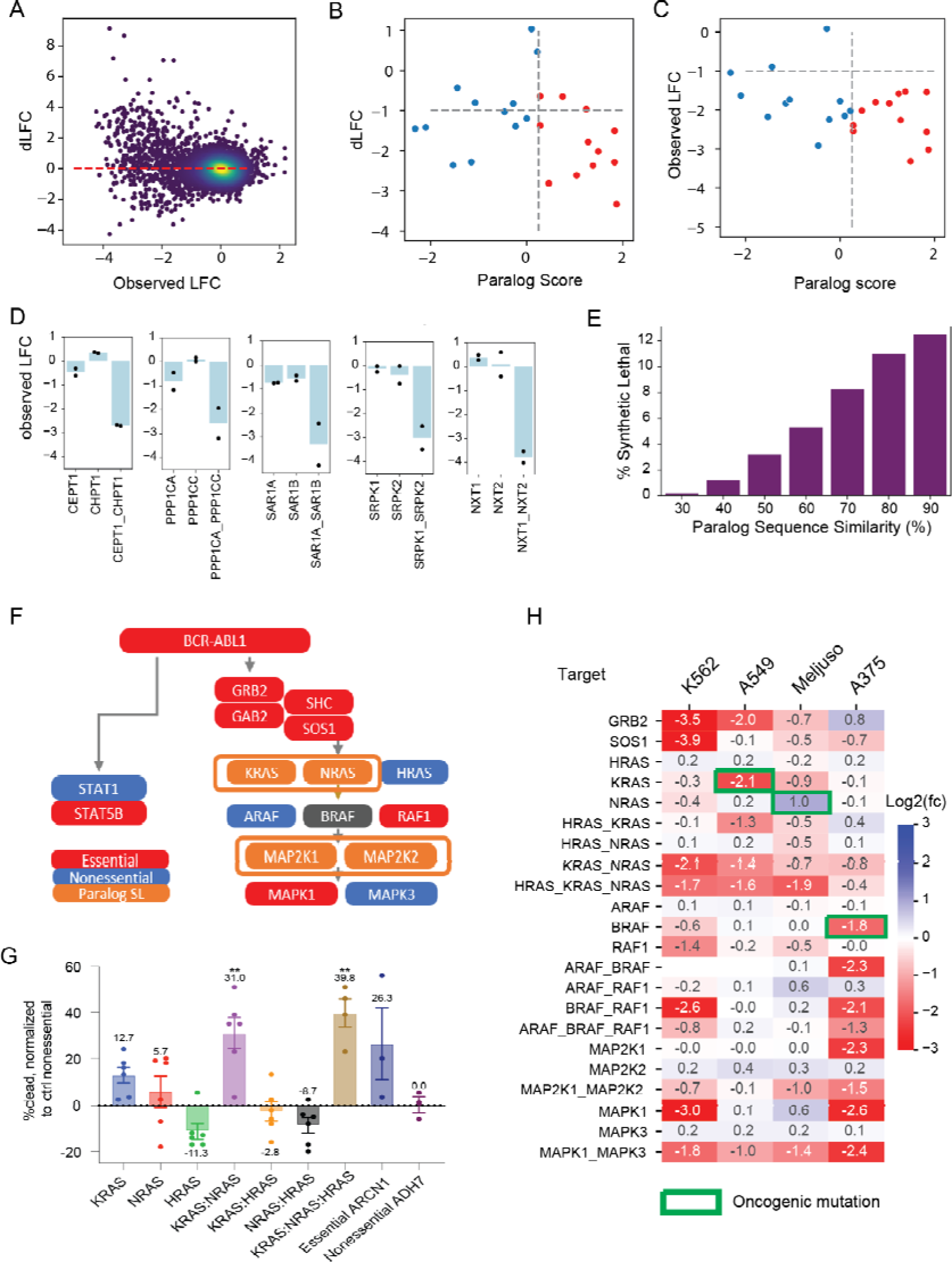
Paralog synthetic lethality with Inzolia. (A) Fold change vs. dLFC for >4,000 paralog families in Meljuso cells. (B) dLFC vs. Paralog Score from meta analysis of published paralog screens. Of 12 paralogs with score > 0.25 (red), 9 show dLFC < -1 in Meljuso cells. C) Fold change vs. paralog score in Meljuso cells. Most scored paralogs are essential, regardless of synthetic lethality. D) Selected synthetic lethals in Meljuso cells showing single and double knockout fitness phenotype. Bar chart, mean fold change. Points indicate fold change of single array of gRNA (mean of 2 replicates). E) Fraction of synthetic lethal paralogs by amino acid sequence similarity in Meljuso cells. F) Pathway activation by BCR-ABL1 fusion in K562 cells. Red, essential gene in in4mer screen; blue, nonessential; orange, synthetic lethal paralog pair. G) Fraction of dead cells, normalized to controls, for single, double, and triple knockouts of RAS genes in K562. KRAS-NRAS joint knockout shows increased cell death. ARCN1, control essential gene. ADH7, control nonessential gene. H) Single, double, and triple knockout phenotype of RTK/MAP kinase pathway genes in all four cell lines. White, target not in library.

The ability to recapitulate known biology is an important control for new technologies, with the MAP kinase pathway a frequently used case study in paralog buffering^21,23^. In K562 cells, the *BCR-ABL* fusion oncogene activates the STAT and MAP kinase pathways, and we classify *ABL1*, *STAT5B*, and the *GRB2/SOS1/GAB2/SHC* signal transduction module as essential genes (Figure 4F). None of the three *RAS* genes are individually essential, but the *KRAS-NRAS* pair shows a strong synthetic lethality. Neither *KRAS-HRAS* nor *HRAS-NRAS* paralogs show genetic interaction, but the three-way *HRAS- KRAS-NRAS* clones also show strong essentiality, almost certainly due to the *KRAS-NRAS* interaction. In an arrayed validation screen, increased cell death after joint KRAS-NRAS knockout confirms this observation (Figure 4G). While it is known that RTK/MAP kinase signal transduction flows through the *RAS* genes, to our knowledge, this is the first time that *KRAS-NRAS* functional buffering has been demonstrated experimentally.

Beyond the *RAS* genes, the rest of the MAP kinase pathway also shows the expected gene essentiality profile in K562 cells (Figure 4H). *RAF1* is strongly essential, and while *BRAF* is slightly below our hit threshold, the *BRAF*-*RAF1* pairwise knockout is consistent with independent additive phenotype. The third member of the paralog family, *ARAF*, is nonessential singly or in combination with the other *RAF* paralogs and has not been shown to operate in this pathway. The MEK kinases, *MAP2K1*/*MAP2K2*, show greater fold change from pairwise loss than from either individually, though below our strict threshold for synthetic lethality. The ERK kinases, *MAPK1*/*MAPK3*, show strong preferential reliance on *MAPK1,* also consistent with DepMap data for K562 cells.

Likewise, the other three cell lines we screened also show oncogene-driven essentiality and GI in the MAPK pathway (Figure 4H). *KRAS* and *BRAF* essentiality in A549 and A375 cells, respectively, are consistent with driver mutations in those genes, though *NRAS* is not detected in *NRAS*-driven Meljuso cells. A375 melanoma cells show isoform dependency on *MAPK1/MAP2K1*, also observed in DepMap screens in *BRAF*-driven melanoma cells. Meljuso cells show interaction between MEK genes and both Meljuso and A549 show GI between ERK genes.

Inzolia screens offer suggestions of polygenic (>2 gene) interactions as well. The library targets 476 paralog triples and quads, and several of these show indications of higher order synthetic lethality (Supplementary Figure 8). We observe an intriguing interaction between *HSPA4* family of Hsp70- related chaperones, where *HSPA4* shows moderate phenotype when knocked out singly or in a pair with either family member, and severe phenotype when all three are targeted. This phenotype is highly variable across the four cell lines. Other candidate background-specific trigenic interactions include the VDAC1/2/3 voltage-gated anion channel family and the ME1/2/3 malic enzyme family, two of which were previously shown to be synthetic lethal in pancreatic cancer under nutrient limiting conditions^32^.

Overall, however, distinguishing three-way synthetic lethal interactions from their composite pairs remains challenging. Even 80% single knockout efficiency can translate into (0.8)^3^ = ∼50% triple knockout efficiency, where cells with incomplete editing can mask severe triple knockout phenotypes.

Higher-order masking interactions, on the other hand, are strikingly visible. Both the core proteasome and the Chaperonin-Containing TCP1 (CCT) complex are composed of several weakly related proteins, which we target with three four-way constructs and numerous two-way constructs. Since both the proteasome and the CCT complex are universally essential to proliferating cells, knockdown of single subunits induces a severe fitness phenotype. Knockout of these genes in pairs or quads yields no additional phenotype, resulting in masking/positive genetic interactions in all four cell lines (Supplementary Figure 8).

Discovering genetic modifiers of drug activity is a common goal of genetic screens. To demonstrate the performance of in4mer constructs in discovering chemogenetic interactions, we screened Meljuso cells in the presence of low-dose MEK inhibitor selumetinib. DrugZ analysis^8^ of single gene targets (Figure 5A) shows that genetic perturbation of the MAP kinase pathway sensitizes cells to the drug (Figure 5A, B). While gene set enrichment analysis^33,34^ identifies subunits of the mitochondrial ribosome, components of the peroxisome, and elements of the Hippo pathway (Figure 5B) as suppressor genes, only two Hippo pathway genes achieved high Z-scores (Figure 5A). DrugZ analysis of pairwise paralog knockouts yielded hits generally consistent with single gene knockout; that is, most paralog knockouts give DrugZ scores consistent with the most extreme single gene knockout (Figure 5C). In some cases, however, combinatorial perturbation of paralogs gave rise to synergistic effects, indicating genetic buffering of chemogenetic interaction. Notably, of the five paralog combinatorial knockouts with Z-scores > 5 and significantly greater effect than their singletons, three encode redundant elements of the Hippo pathway, including *STK3/STK4, LATS1/LATS2*, and *MOB1A/MOB1B* (Figure 5C,D).

**Figure 5.**
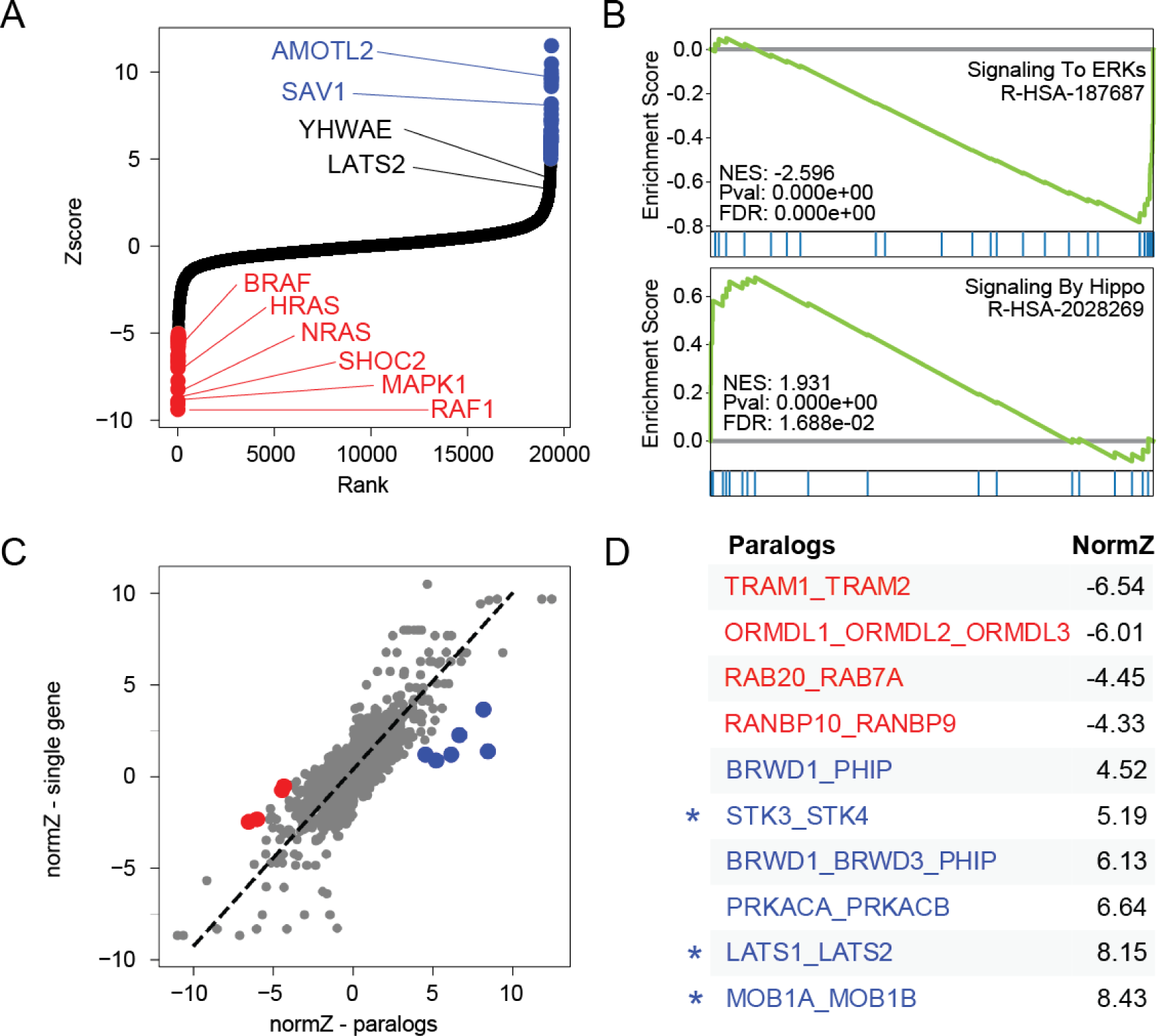
Synthetic chemogenetic interactions. Meljuso cells were cultured in the presence of MEK inhibitor selumetinib and screened for chemogenetic interactions (A) DrugZ scores of single gene knockouts. Selected genes in the MAPK and Hippo pathways highlighted. (B) Selected GSEA results for gene sets conferring sensitivity (ERK signaling) or resistance (Hippo signaling) to MEKi. (C) Comparing DrugZ scores of paralogs (x-axis) vs. the most extreme Z score of the single gene knockout (y-axis) shows that most pairwise perturbagens yield similar phenotype as singletons. Outliers in red (synergistic) or blue (suppressor). (D) Synergistic and suppressor paralog knockouts from (C). Asterisk indicates functional buffering in the Hippo pathway, masking phenotype in monogenic knockout screens.

In total, the Inzolia library includes ∼50k unique guide arrays, with ∼40k targeting single genes and 9,822 arrays targeting paralog doubles, triples, and quads. Inzolia is therefore on par with latest-generation genome-scale CRISPR/Cas knockout libraries^26,35–38^ (Figure 6), and is unique among such libraries in including thousands of reagents targeting paralogs. Moreover, the efficiency gain realized by having two guides targeting each of two genes in a paralog pair makes detection of genetic interactions tractable with only six reagents per gene pair, a fivefold improvement over the prior state of the art (Figure 6).

**Figure 6.**
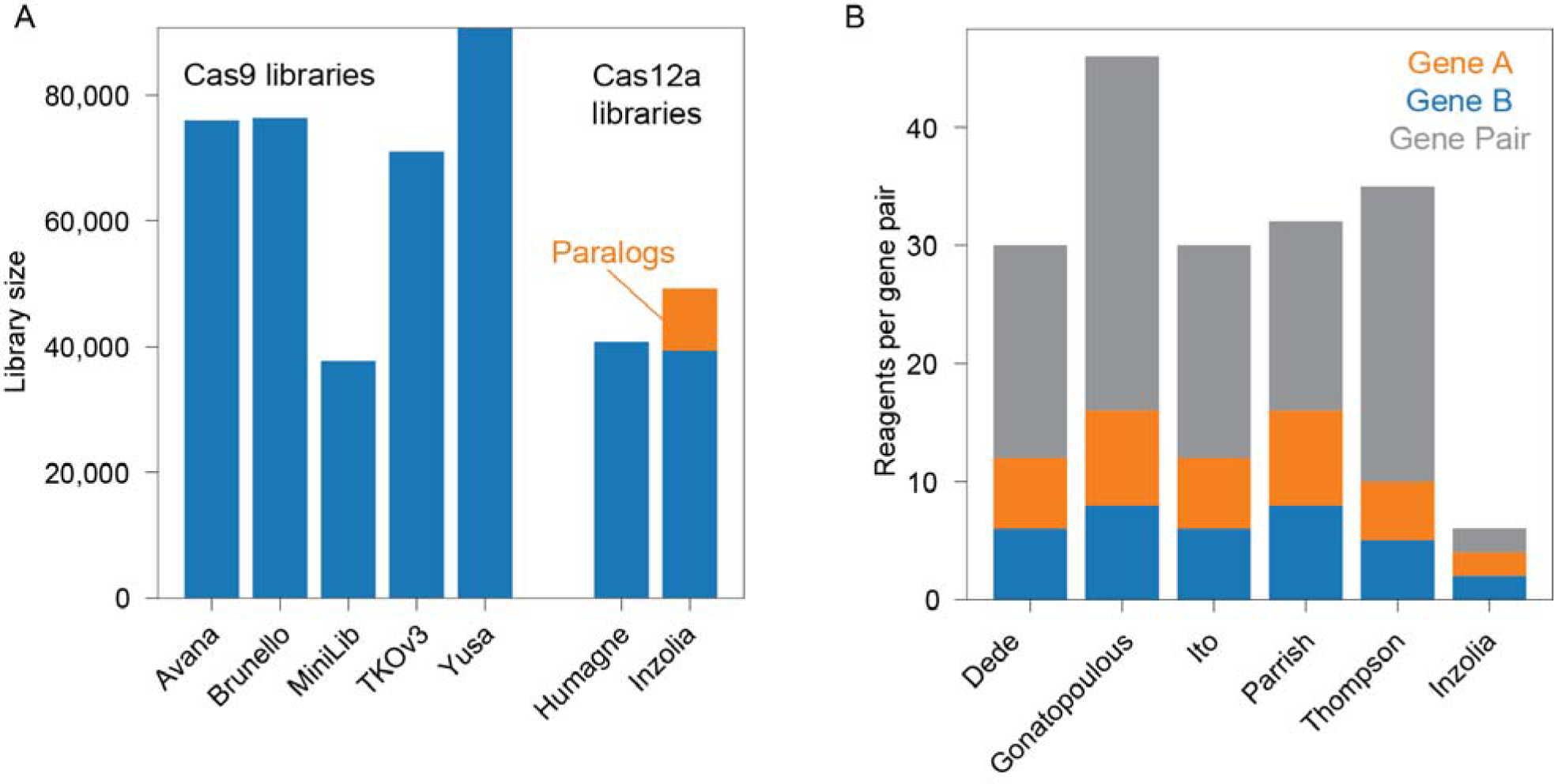
Library size comparison. (A) Representative Cas9 and Cas12a whole-genome libraries. Inzolia library targets 19k protein coding genes and additionally includes 9,822 guide arrays targeting paralog doubles, triples, and quads. (B) Five recent publications screening for genetic interactions between paralogs. Bar plot shows number of reagents per paralog pair tested, including single and double knockouts.

## Conclusions

The rapid ascendancy of CRISPR-mediated genetic perturbation technologies over RNA interference methods was driven by major advances in assay sensitivity and specificity, with the absence of established gold standards arguably contributing to the shortcomings of RNAi-based studies of mammalian gene function^39^. We^38,40^ and others^2^ have created widely used reference sets of essential and nonessential genes for use in quality control of monogenic loss of fitness screens. As CRISPR perturbation technology has advanced into genetic interactions, it has become clear that a similar gold standard for synthetic lethals is needed^30^. Our meta-analysis of published screens for paralog synthetic lethals in human cells shows wide divergence in the paralogs assayed by each study and in the repeatability of each screen, as measured by the Jaccard coefficient of hits in different cell lines. We reasoned that paralogs that showed synthetic lethality within and across screening platforms are likely to be globally synthetic lethal, analogous to core essential genes, and the fact that 12 of our 13 candidate reference paralogs show more than 70% identity (and all are constitutively expressed) is consistent with this interpretation.

Notably, the engineered Cas12a endonuclease, developed in Kleinstiver et al.^25^ and deployed in combinatorial screens in DeWeirdt et al. and paralog screens in Dede et al.^18^ performed markedly better in terms of replicability. Based on this and our prior work with the CRISPR/Cas12a screens^26,27^, we tested the limits of the Cas12a system expressing guide arrays from the Pol III U6 promoter in a custom one-component lentiviral vector, pRDA_550. For longer arrays of seven independent gRNA, we observed that position-specific loss of knockout efficiency did not arise until after the fourth or fifth gRNA in the array. From this we developed the in4mer platform for arrays encoding four independent gRNA, each with an optimized spacer sequence from the CRISPick algorithm and with diverse but proven DR sequences to minimize the chance of recombination. By targeting single genes with four independent gRNA, we lower the odds that any single guide fails to induce the desired phenotype, extending the development of the Humagne library in work from DeWeirdt et al^26^. As with the Humagne library, having multiple independent gRNA on each array reduces the total number of reagents required to induce reliable gene perturbation.

To construct our Inzolia genome-scale human library, we began with reagents targeting single protein-coding genes and added arrays targeting more than four thousand paralog pairs, triples, and quads, with the in4mer arrays encoding two guides targeting each of the two genes in a pair or one guide per gene in a triple or quad family. Inzolia screens show high (at least 75%) sensitivity to detect synthetic lethals with just two reagents targeting each single knockout and two reagents targeting the double knockout, and offer the potential for novel biology arising from three- and four-way paralog synthetic lethals. The Inzolia library is thus a smaller and more efficient whole-genome library that addresses one of the major gaps of monogenic perturbation libraries: functional buffering by paralogs.

Beyond the paralogs in the Inzolia library, the in4mer system is a highly customizable platform offering a significant advance in the study of genetic interactions. Compared to the five paralog synthetic lethal studies, with each using at least thirty constructs per gene pair tested, in4mer requires fivefold fewer reagents for the same assay. This improvement has major implications for the cost effectiveness in genetic interaction assays in mammalian cells, where the number of gene pairs and the diversity of cell/tumor lineages and genotypes combine to yield a vast search space. A fivefold reduction in experimental footprint could offer a correspondingly larger search space for the same effort, or the same search space across more backgrounds (e.g. cell lines) or environments (e.g. chemogenetic interactions) at nearly the same cost as a single screen with an equivalent combinatorial Cas9 system, with the Cas12a system yielding greater sensitivity and robustness. Moreover, custom library construction leverages the other key advantage of the Cas12a system: each library is constructed from a single ∼200mer oligo pool and both cloning and amplicon sequencing are performed using essentially the same protocols as single-guide Cas9 screening, albeit with longer sequencing reads. With the in4mer system, a wider swath of the research community will be able to add targeted genetic interaction surveys to their experimental toolkits.

## Acknowledgments

NEA, LLW, XM, MC and TH were supported by NIGMS grant R35GM130119, NCI grant U01CA275886, and CPRIT grant RP210173. NEA and SHW are supported by National Institutes of Health grant R01GM139980. TH is a CPRIT Scholar in Cancer Research and an Andrew Sabin Family Fellow. CL is an Odyssey Fellow and is supported by the Odyssey Program and Odyssey Expansion Fund at MD Anderson. This work was additionally supported by the NCI Cancer Center Support Grant P30CA16672 (TH) and NCI Cancer Moonshot Initiative U01CA250565 (JGD).

## COI

Disclosure Statement: JGD consults for Microsoft Research, Abata Therapeutics, Servier, Maze Therapeutics, BioNTech, Sangamo, and Pfizer. JGD consults for and has equity in Tango Therapeutics. JGD serves as a paid scientific advisor to the Laboratory for Genomics Research, funded in part by GlaxoSmithKline. JGD receives funding support from the Functional Genomics Consortium: Abbvie, Bristol Myers Squibb, Janssen, Merck, and Vir Biotechnology. JGD’s interests were reviewed and are managed by the Broad Institute in accordance with its conflict of interest policies.

## Methods

### Paralog meta-analysis

#### Data preprocessing

To reanalyze the data from the 5 paralog screens, raw read counts were downloaded, and the same pipeline was applied to all of them. A pseudocount of 5 reads was added to each construct in each replicate, and total read counts were normalized to 500 reads per construct. Log2 fold change (LFC) for each guide at late time point was calculated relative to the plasmid sequence counts.

The data from each study (except Thompson) were divided into three groups; the constructs that target single genes paired with non-essential/non-targeting gRNAs (N) in the first position (gene_N), in the second position (N_gene) and constructs that target gene pairs (A_B). LFC values of each group were scaled individually so that the mode of each group was set to zero. Next, all three groups were merged in one table. Before dividing Ito’s dataset into three groups, LFC values were scaled such that the mode of negative controls (non-essential_AAVS1) would be zero and also TRIM family was removed from this dataset to avoid false paralog pair discovery ^13^. Since in Thompson’s study there was just one position for singleton constructs, LFC values were scaled so that the mode of negative controls (non- essential_Fluc) was set to zero. In the next step, LFC of each construct was calculated by the mean of LFC across different replicates.

#### Calculating genetic interaction

For each gene, single gene mutant fitness (SMF) was calculated as the mean construct log fold change of gene-control constructs. The control was either non-essential genes or non-targeting gRNAs. For each gene pair, the expected double mutant fitness (DMF) of genes 1 and 2 was calculated as the sum of SMF of gene 1 and SMF of gene 2. The difference between expected and observed DMF, the mean LFC of all constructs targeting genes 1 and 2, was called dLFC.

Next step was calculating a modified Cohen’s D between observed and expected distribution of LFC of gRNAs targeting genes. Expected distribution of gRNAs targeting a gene pair, was calculated using expected mean and expected standard deviation.

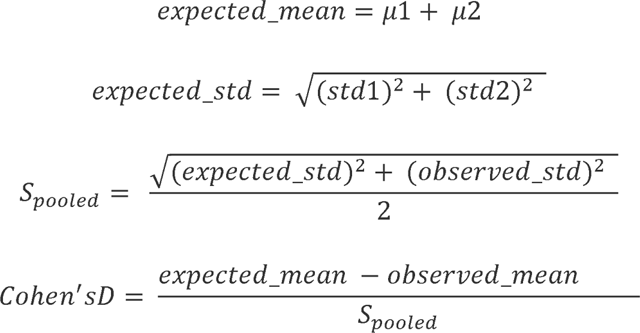

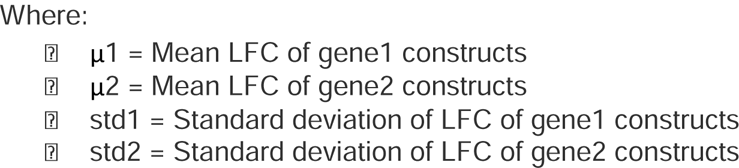

In each cell line, the paralog pairs with dLFC < -1 and Cohen’s D > 0.8 were selected as hits. Cohen’s D more than 0.8 indicates large effect size between two groups, meaning that our expected and observed distribution of gRNAs are meaningfully separated. In total 388 paralog pairs were identified as hits across all the studies.

To identify the most consistent method in terms of hit identification, the Jaccard similarity coefficient of every pair of cell lines in each study was calculated by taking the ratio of intersection of hits over union of hits. For the studies that screened more than two cell lines, the final platform weight was the median of the calculated Jaccard coefficients of all pairs of cell lines.

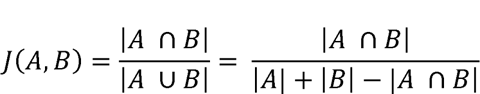

#### Scoring Paralog Pairs

Each hit was scored based on the cell lines in which it was identified as a hit; cell lines were weighted based on the platform weight described above. We defined the “paralog score” as the sum of platform weights of cell lines in which the paralog pair was identified as a hit minus the sum of platform weights of cell lines in which the paralog pair was assayed but not identified as a hit (a “miss”). The distribution of scores is shown in Figure 1. Gene pairs with paralog score > 0.25 and were identified as a hit in two or more studies were listed as candidate gold standard paralog synthetic lethals.

### Combinatorial CRISPR screen design

#### One-component CRISPR/enCas12a vector

To construct an all-in-one vector for expression of both Cas12a and a guide array, we first swapped in puromycin resistance in place of blasticidin resistance from pRDA_174 (Addgene 136476). We then tested four locations for the insertion of a U6-guide expression cassette; notably this cassette needs to be oriented in the opposite direction of the primary lentiviral transcript to prevent Cas12a-mediated processing during viral packaging in 293T cells. The construct with the best-performing location, between the cPPT and the EF-1α promoter, was designed pRDA_550 (Addgene #203398). Synthesis of DNA and custom cloning was performed by Genscript.

#### 7Mer library production

An oligonucleotide pool consisting of 7 Essential and 7 Non-Essential gene crRNAs with their nearby DR, BsmBI recognition as well as overhang sequence was synthesized by Integrated DNA Technologies. The pool was amplified by asymmetric PCR followed by being assembled into PRDA_550 vector to acquire the designed library through NEBridge® Golden Gate Assembly Kit (BsmBI-v2) (New England Biolabs). The assembled product was transformed into NEB® Stable Competent E. coli (High Efficiency) cells and the plasmid DNA was purified using the PureLink™ Plasmid Purification Kit (Invitrogen). Three oligonucleotide pools were cloned separately and pooled together to acquire the final 7mer library. The library was sequenced to confirm uniform and complete library representation.

#### Paralog selection for IN4MER/INZOLIA

Human paralogs and percent identity data were imported from BioMart, which reports both AB and BA percent identity (these can differ if the two genes encode proteins of different lengths) Mean percent identity ((AB + BA) / 2)and delta percent identity (|AB - BA|) between paralogs were then calculated, and for the prototype library, paralogs with mean percent identity between 30% and 99% and delta percent identity < 10% were selected (Supplementary Figure 5). Next, CCLE expression data was downloaded, and the mean and standard deviation of expression across all CCLE samples was calculated for each gene. Paralogs where both genes had mean expression > 2 and stdev < 1.5 were selected (i.e. constitutively expressed genes).

Finally, to identify and include paralog families of size > 2, we applied a “difference from top paralog” filter. For each gene A in the pool, we identified its top paralog B by max sequence identity. Then for each other candidate paralog C, we calculated the drop in sequence identity, AB - AC (see distribution of drop % in Supplementary Figure 5). For the prototype library, we defined A,B,C as being in the same family if AB - AC < 10%.

For the final Inzolia library, we relaxed several of these filters. The delta percent identity filter and the expression variance filter were removed entirely, and the difference from top paralog filter was expanded to 20%. The mean expression filter was retained. These three filtering steps resulted in a total of 4435 paralog pairs included in the Inzolia pool library.

#### IN4MER Prototype library production (MD Anderson)

Oligonucleotide pools consisting of designed four-plex guide arrays were synthesized by Twist Bioscience. The prototype pool consists of 43,972 arrays targeting 19,687 single genes, 2,082 paralog pairs, 167 paralog triples, and 48 paralog quads.

5’-*AATGATACGGCGACCACCGA***cgtctcgAGAT**nnnnnnnnnnnnnnnnnnnnTAATTTCTACTATTGTAGATn nnnnnnnnnnnnnnnnnnnAAATTTCTACTCTAGTAGATnnnnnnnnnnnnnnnnnnnnTAATTTCTACTGTCGT AGATnnnnnnnnnnnnnnnnnnnnTTTTTT**GAATggagacg***ATCTCGTATGCCGTCTTCTGCTTG*-3’.

*ITALIC* - primer sequence

**BOLD** - BsmBI restriction sequence. Overhang in CAPS. nnnnn - guide sequence

UNDERLINED - DR sequence The pool of guide arrays was PCR amplified using KAPA HiFi 2X HotStart ReadyMix (Roche) using 20 ng of starting template per 25 μl reaction using forward and reverse primers:

FWD (5’-AATGATACGGCGACCACCGA-3’) REV (5’-CAAGCAGAAGACGGCATACGAGAT-3’) at a final concentration of 0.3 μM and the following conditions: denaturation at 95°C for 3 min, followed by 12 cycles of 20 s at 98°C, 30 s at 60°C, and 30 s at 72°C, followed by a final extension of 1 min at 72°C. The resulting amplicon was purified by the Monarch PCR & DNA Cleanup Kit (New England Biolabs) and cloned into the pRDA-550 vector by NEBridge® Golden Gate Assembly Kit (BsmBI-v2) The product from assembly reaction was purified and electroporated into Endura Electrocompetent cells (Lucigen). Transformed bacteria were diluted 1:100 in 2xYT medium containing 100 μg/ml carbenicillin (Sigma) and grown at 30 °C for 16 hours. The plasmid DNA was extracted by PureLink™ Plasmid Purification Kit (Invitrogen). The library was sequenced to confirm uniform and complete library representation.

#### INZOLIA library production (Broad Institute)

The final Inzolia pool consists of arrays targeting 19,687 single genes, 4,435 paralog pairs, 376 paralog triples, and 100 paralog quads, plus 20 arrays targeting EGFP, 500 targeting intergenic loci, and 50 encoding nontargeting guides. Each array in the oligonucleotide pools is constructed as follows:

5’-*AGGCACTTGCTCGTACGACG***cgtctcgAGAT**nnnnnnnnnnnnnnnnnnnnTAATTTCTACTATTGTAGATn nnnnnnnnnnnnnnnnnnnAAATTTCTACTCTAGTAGATnnnnnnnnnnnnnnnnnnnnTAATTTCTACTGTCGT AGATnnnnnnnnnnnnnnnnnnnnTTTTTT**GAATggagacg***TTAAGGTGCCGGGCCCACAT*-3’.

*ITALIC* - primer sequence

**BOLD** - BsmBI restriction sequence. Overhang in CAPS. nnnnn - guide sequence

UNDERLINED - DR sequence

The pool of guide arrays was PCR amplified using NEBNext® High-Fidelity 2X PCR Master Mix (NEB) using 196 ng of starting template per 50 μl reaction using forward and reverse primers:

FWD (5’-*AGGCACTTGCTCGTACGACG*-3’) REV (5’-ATGTGGGCCCGGCACCTTAA-3’) at a final concentration of 0.5 μM and the following conditions: denaturation at 98°C for 1 min, followed by 7 cycles of 30 s at 98°C, 30 s at 53°C, and 30 s at 72°C, followed by a final extension of 5 min at 72°C. The resulting amplicon was purified by the Qiaquick PCR Purification Kit (Qiagen) and cloned into the pRDA-550 and pRDA-052 via Golden Gate cloning with Esp3I (Fisher Scientific) and T7 ligase (Epizyme). The assembly product was purified by isopropanol precipitation, electroporated into Stbl4 electrocompetent cells (Life Technologies) and grown at 37 °C for 16 hours on agar with 100 ug/mL carbenicillin. Colonies were scraped and plasmid DNA (pDNA) was extracted via HiSpeed Plasmid Maxi (Qiagen). The library was sequenced to confirm uniform and complete library representation.

#### Cell culture

K562 and A549 cells were a gift from Tim Heffernan. A375 and MelJuso were obtained from the Cancer Cell Line Encyclopedia. Cell line identities were confirmed by STR fingerprinting by M.D. Anderson Cancer Center’s Cytogenetic and Cell Authentication Core. All cell lines were routinely tested for mycoplasma contamination using cells cultured in non-antibiotic medium (PlasmoTest Mycoplasma Detection Assay, InvivoGen).

All cell lines were grown at 37°C in humidified incubators at 5.0% CO_2_ and passaged to maintain exponential growth.

For each cell line, the following medium and concentration of polybrene (EMD Millipore) and puromycin (Gibco) were used: K562: RPMI+10% FBS, 8 μg/ml, 2 μg/ml A549: DMEM+10%FBS, 8 μg/ml, 2 μg/ml A375: RPMI+10% FBS, 1 μg/ml, 1 μg/ml MelJuso: RPMI+10% FBS, 4 μg/ml, 1 μg/ml

#### Cas12a Screens

Lentivirus was produced by the University of Michigan Vector Core (prototype) or the Broad GPP (Inzolia). Virus stocks were not titered in advance. Transduction of the cells was performed at 1X concentration of virus with corresponding polybrene.

Non-transduced cells were eliminated via selection puromycin dihydrochloride. Selection was maintained until all non-transduced control cells reached 0% viability. Once selection with puromycin was complete, surviving cells were pooled and 500x coverage cells were harvested for a T0 sample. After T0, cells were harvested at 500X coverage on corresponding days.

#### Prototype In4mer library Genomic DNA preparation and sequencing (MD Anderson)

Genomic DNA (gDNA) was extracted using the Mag-Bind® Blood & Tissue DNA HDQ 96 Kit (Omega Bio-tek) and quantified by the Qubit™ dsDNA Quantification Assay Kits (ThermoFisher).

Illumina-compatible guide array amplicons were generated by amplification of the gDNA in a one-step PCR. Indexed PCR primers were synthesized by Integrated DNA Technologies using the standard 8nt indexes from Illumina (D501-D508 and D701-D712) as follows:

forward primer:5’-**AATGATACGGCGACCACCGAGATCTACAC**nnnnnnnnACACTCTTTCCCTACACGACGCTCTTCCGATCT*CTTGTGGAAAGGACGAAACACCG*-3’ (**i5 flow cell adapter** – i5 index – i5 read1 primer binding site – *Amplicon annealing sequence*)

reverse primer:

5’-**CAAGCAGAAGACGGCATACGAGAT**nnnnnnnnGTGACTGGAGTTCAGACGTGTGCTCTTCCGATCT *ACCGACTCGGTGCCACTTTTTCAAGACCAG*-3’ (**i7 flow cell adapter** – i7 index – i7 read2 primer binding site – *Amplicon annealing sequence*).

At least ∼200X coverage gDNA per replicate across multiple reactions were amplified. Each gDNA sample was first divided into multiple 50 μl reactions with most 2.5ug gDNA per reaction. Each reaction contained 1ul each primer (10 μM), 1 μl 50X dNTPs, 5% DMSO, 5 μl 10X Titanium Taq Buffer, and 1 μl 10X Titanium Taq DNA Polymerase (Takara). The PCR conditions were: denaturation at 95 C for 60 s, followed by 25 cycles of 30 s at 95°C and 1 min at 68°C, followed by a final extension at 68°C for 3 min. After the PCR, all reactions from the same sample were pooled and then purified by E-Gel™ SizeSelect™ II Agarose Gels, 2% (ThermoFisher). Purified amplicons were quantified by Qubit™ dsDNA Quantification Assay Kits (ThermoFisher) and validated by D1000 ScreenTape Assay for TapeStation Systems (Agilent) (360bp for IN4MER, 501bp for 7Mer). Purified amplicons were then pooled (with 30% customized random library to increase the diversity) and sequencing was performed by NextSeq 500 sequencing platform (Illumina) with custom primers (Integrated DNA Technologies):

forward:5’-CTTGTGGAAAGGACGAAACACCGGTAATTTCTACTCTTGTAGAT-3’ reverse: 5’-ACCGACTCGGTGCCACTTTTTCAAGACCAG-3’ (7Mer only).

The IN4MER library was sequenced by read format of 151-8-8, single-end and the 7Mer library was sequenced by read format of 151-8-8-151, paired-end.

#### Inzolia Genomic DNA preparation and sequencing (Broad Institute)

Genomic DNA (gDNA) was extracted using the Mag-Bind® Blood & Tissue DNA HDQ 96 Kit (Omega Bio-tek) and quantified by the Qubit™ dsDNA Quantification Assay Kits (ThermoFisher).

Illumina-compatible guide array amplicons were generated by amplification of the gDNA in a one-step PCR. Indexed PCR primers were synthesized by Integrated DNA Technologies using the standard 8nt indexes from Illumina (D501-D508 and D701-D712). The sequences for the primer sets were used for the libraries cloned into pRDA-052 or pRDA-550 are listed below:

pRDA-052 compatible forward primer:

5’-**AATGATACGGCGACCACCGAGATCTACAC**nnnnnnnnACACTCTTTCCCTACACGACGCTCTTCCG

ATCT*TGTGGAAAGGACGAAACACCG*-3’ (**i5 flow cell adapter** – i5 index – i5 read1 primer binding site – [stagger region of 0-8 nt] – *Amplicon annealing sequence*)

pRDA-052 compatible reverse primer:

5’-**CAAGCAGAAGACGGCATACGAGAT**nnnnnnnnGTGACTGGAGTTCAGACGTGTGCTCTTCCGATCT

[s]*TCTACTATTCTTTCCCCTGCACTGT* -3’ (**i7 flow cell adapter** – i7 index – i7 read2 primer binding site – [stagger region of 0-8 nt] – *Amplicon annealing sequence*).

pRDA-550 compatible forward primer:

5’-**AATGATACGGCGACCACCGAGATCTACAC**nnnnnnnnACACTCTTTCCCTACACGACGCTCTTCCG

ATCT*TGTGGAAAGGACGAAACACCG*-3’ (**i5 flow cell adapter** – i5 index – i5 read1 primer binding site – [stagger region of 0-8 nt] – *Amplicon annealing sequence*)

pRDA-550 compatible reverse primer:

5’-**CAAGCAGAAGACGGCATACGAGAT**nnnnnnnnGTGACTGGAGTTCAGACGTGTGCTCTTCCGATCT

[s]*ACCGACTCGGTGCCACTTTTTCAA*-3’ (**i7 flow cell adapter** – i7 index – i7 read2 primer binding site – [stagger region of 0-8 nt] – *Amplicon annealing sequence*).

At least ∼200X coverage gDNA per replicate across multiple reactions were amplified. Each gDNA sample was first divided into multiple 100 μl reactions with most 10 μg gDNA per reaction. Each reaction contained 0.5 μl forward primer (100 μM), 10uL reverse primer (5 uM) 8 μl dNTPs, 5 μl DMSO, 10 μl 10X Titanium Taq Buffer, and 1.5μl Titanium Taq DNA Polymerase (Takara). The PCR conditions were: An initial denaturation at 95°C for 60 s, followed by 28 cycles of 30 s at 94°C, 30 s at 52°C, and 30 s at 72 °C followed by a final extension at 72°C for 10 min. After the PCR, all reactions from the same sample were pooled and purified with Agencourt AMPure XP SPRI beads according to the manufacturer protocol (Beckman Coulter). Purified amplicons were quantified by Qubit™ dsDNA Quantification Assay Kits (ThermoFisher) and sequenced on a HiSeq2500 with a Rapid Run (200 cycle) kit (Illumina).

#### 7Mer screen data analysis

Reads for each reagent were counted using only exact matches to the entire 281 nucleotide 7mer sequence, excluding the leading DR (7 23mer spacer sequences + 6 20mer DR sequences). Fold changes were calculated relative to the mean of the T0 samples, and averaged across replicates. For each sample (T7/14/21), fold changes were normalized by subtracting the mean fold change of arrays with 7 nonessentials; i.e. setting no-essentials guides to zero.

We expected that the selected essential genes would not show any pairwise or higher order interactions, and thus should be governed by the multiplicative model of genetic interaction. To evaluate this model, we fit a regression model:

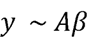

where A is a binary matrix of 7mer guide arrays (rows, k=384) by positions (columns, n=7), with A_i,j_ = 1 if guide array i targets an essential gene at position *j* and 0 if not. y is the vector of normalized observed fold changes, and the n-length vector f3 coefficients represent the single gene knockout phenotype learned from the model. We filtered this construct for reagents that encoded two or fewer essential genes (k=87 rows). After linear fit, we compared the predicted zero, one, and two gene knockout fitness profiles (by summing the f3 coefficients for each gene) to the mean observed knockout fitness. R^2^ values for each pool ranged from 0.78 to 0.91, and the overall quality of the linear fit supports the multiplicative model for non-interacting genes as assayed by combinatorial CRISPR knockouts of up to two genes. An accurate null model for noninteraction is critical for detecting and classifying deviations from this model that reflect positive or negative genetic interactions.

#### IN4MER/Inzolia screen data analysis

In4mer library sequencing reads were mapped to the library using only perfect matches. BAGEL2 was used to normalize sample level read counts and to calculate fold changes relative to the T0 reference using the BAGEL2.py *fc* option with default parameters^41^. Essential and nonessential genes were defined using the Hart reference sets from^38,40^. Since the library targets both individual genes and specific gene sets (paralogs), we calculated the average gene/gene set (hereafter ‘gene’) log fold change as the mean of the clone-level fold changes across two replicates. All fold changes are calculated in log2 space. Cohen’s D statistics were calculated in Python as described in Paralog meta-analysis above. Data for recall-precision curves were calculated using BAGEL2. We set an arbitrary threshold of fc < -1 for essential genes.

For genetic interaction analysis, the expected fold change was calculated as the sum of the gene-level fold changes for each individual gene in the gene set. Expected fc was subtracted from observed fc to calculate delta log fold change, dLFC, where negative dLFC indicates synthetic/synergistic interactions with more severe negative phenotype, and positive dLFC indicates positive/suppressor/masking interactions with less severe negative or more positive phenotype than expected. We set an arbitrary threshold of dLFC < -1 for synthetic lethality, and > +1 for masking/suppressor interactions.

#### RAS synthetic lethal validation

An arrayed knockout apoptosis assay approach was adopted to validate RAS synthetic lethality in K562. Two guides were selected for each of the three RAS genes, and two clones were designed for each target/gene combination. Guide RNAs were selected through CRISPick and gblocks (same construct as Inzolia library) were synthesized by Integrated DNA Technologies. The arrays were individually cloned into the pRDA_550 backbone and plasmids were validated by Sanger sequencing. The plasmids were then individually transfected to K562 cells via the Neon Transfection System (Invitrogen). Each group was transfected with 2 μg of DNA per 2 × 10^6^ cells, using the recommended setting for K562 electroporation with one pulse at 1000 v, 50 ms. Non-transfected cells were eliminated through puromycin selection, which was maintained until non-transfected control cells reached 0% viability. Triplicate wells were maintained after selection until the end of the experiment. Cell viability, total cell numbers, live cell size and dead cell size data were collected through reading Trypan Blue (Gibco) stained cells via Countess II FL (Thermo Fisher) at each passage until 9 days after puromycin selection, in line with Inzolia screen end point of 8 days in K562 cells. Percent dead cells were normalized to negative control and unpaired T tests were conducted to compare experimental groups against the negative control for statistical significance.

## Notes

### Summary of Updates

Significant additional data and updated version of the genome-scale library, "Inzolia"

